# Personalized Risk Prediction for Type 2 Diabetes: the Potential of Genetic Risk Scores

**DOI:** 10.1101/041731

**Authors:** Kristi Läll, Reedik Mägi, Andrew Morris, Andres Metspalu, Krista Fischer

## Abstract

**Purpose:** The study aims to develop a Genetic Risk Score (GRS) for the prediction of Type 2 Diabetes (T2D) that could be used for risk assessment in general population.

**Methods:** Using the results of genome-wide association studies, we develop a *doubly-weighted* GRS for the prediction of T2D risk, aiming to capture the effect of 1000 single nucleotide polymorphisms. The GRS is evaluated in the Estonian Biobank cohort (n=10273), analysing its effect on prevalent and incident T2D, while adjusting for other predictors. We assessed the effect of GRS on all-cause and cardiovascular mortality and its association with other T2D risk factors, and conducted the reclassification analysis.

**Results:** The adjusted hazard for incident T2D is 1.90 (95% CI 1.48, 2.44) times higher and for cardiovascular mortality 1.27 (95% CI 1.10, 1.46) times higher in the highest GRS quintile compared to the rest of the cohort. No significant association between BMI and GRS is found in T2D-free individuals. Adding GRS to the prediction model for 5-year T2D risks results in continuous Net Reclassification Improvement of 0.26 (95% CI 0.15, 0.38).

**Conclusion:** The proposed GRS would considerably improve the accuracy of T2D risk prediction when added to the set of predictors used so far.

## INTRODUCTION

The increasing prevalence of Type 2 Diabetes (T2D) is currently one of the greatest challenges for public health, in both developed and developing countries alike. In 2012, 371 million people - approximately 8.3% of the world’s adult population - were estimated to be living with diabetes^1^. Diabetes is a leading cause of cardiovascular disease, renal disease, blindness, and limb amputation. T2D, accounting for 80-90% of all diabetes in Europe, decreases life expectancy by 5-10 years^2^.

As the onset of T2D can be postponed or partially prevented by changes in the lifestyles of high-risk subjects^3^, cost-effectiveness of lifestyle (and other) interventions can be increased by improving the precision of risk prediction, thereby enabling targeting of the individuals at highest risk.

Although obesity is the strongest predictor of T2D, it is also known that heritability of T2D is 26%-69%, depending on age of onset^4,5^, motivating the search for genetic predictors for T2D. However, despite the large number of published genome-wide association studies (GWAS) of T2D so far, there is still some scepticism on the practical value of identified single nucleotide polymorphisms (SNPs) in personalized risk prediction for the disease. The main reason is that the effect of individual SNPs on complex common disease phenotypes is relatively weak and/or adds little to predictions based on lifestyle, demographic and clinical factors^6,7^.

In GWAS, SNPs need to meet the stringent genome-wide threshold, usually set to p = 5*10^−8^, to be significantly associated with the trait. Even though the sample sizes in GWAS have been increasing steadily over the years, they are still insufficient for SNPs with small effects to pass that threshold^8^. This could explain why it has been shown that all common variants across the genome actually explain much higher proportion of heritability (50% or more) in many complex traits than one could see based on a small subset of significant SNPs only^9,10^.

To actually explain a meaningful proportion of variability in a complex trait and, more importantly, to use this knowledge in risk assessment at an individual level, one needs to construct a numeric measure with acceptable predictive power – a genetic (polygenic) risk score (GRS) based on a large number of genotyped variants. Our aim is to develop a GRS that can be implemented in routine personalized risk prediction to improve T2D risk stratification in the general population. For that purpose, we will use two sources of data: firstly, results of large-scale meta-analyses of GWAS to obtain effect estimates for individual SNPs with best possible prediction^11^. Secondly, individual level data of a relatively large validation cohort from the Estonian Biobank is used to compare different versions of GRS and decide on applicability of the best GRS in practical risk prediction. The best-fitting GRS for prevalent cases is then further validated in the analysis of incident T2D patients, obtained by linking the Estonian Biobank cohort database to electronic health records of the participants.

## MATERIALS AND METHODS

### Estonian Biobank Cohort and Genotyping

The Estonian Biobank (Estonian Genome Center, University of Tartu) was established in 2002, with the long-term purpose of implementation of research results to public health and medicine in Estonia. Between 2002 and 2011, the Estonian Biobank has recruited a cohort of 51380 participants which includes adults from all counties of Estonia and accounts for approximately 5% of the Estonian adult population during the recruitment period. A broad informed consent signed by participants enables the use of the data for various health research purposes, as well as linkage of the data with other health-related databases and registries. An extensive phenotype questionnaire and measurement panel, together with follow-up data from linkage with national health-related registries and electronic health records (Estonian Health Insurance database), allows assessment of the effects of classical epidemiological risk factors on the incidence of common complex diseases, such as T2D.

In the present study, a genotyped subset of 10273 individuals (including 1181 prevalent T2D cases) from the cohort has been analysed. The DNA samples of this subset are genotyped using either the Illumina Human OmniExpress (a random sample of 8085 individuals) or Illumina Cardio-MetaboChip (a case-control sample of 942 T2D cases, 680 cases of Coronary Artery Disease and 903 random controls) genome-wide arrays. For 337 individuals (including 169 T2D cases) genotyped by both arrays, the genotype data from Cardio-MetaboChip array was used.

During the average follow-up time of 5.63 years, 386 incident T2D cases were observed (in individuals free of T2D at recruitment) by 1 April 2014. Moreover, a total of 1994 individuals of the analysed set had died by 1 September 2015 (including 1069 deaths due to cardiovascular causes).

The baseline phenotype data (Table 1) used for this study consists of age, gender, BMI and prevalent T2D status. For a subset (*n* = 6064), data on plasma glucose level, as well as lipoprotein profiles (LDL and HDL cholesterol, triglycerides and total cholesterol) obtained by Nuclear Magnetic Resonance (NMR) profiling is available (non-fasting measurements, with information on the time of last meal available for adjustment).

**Table 1.**
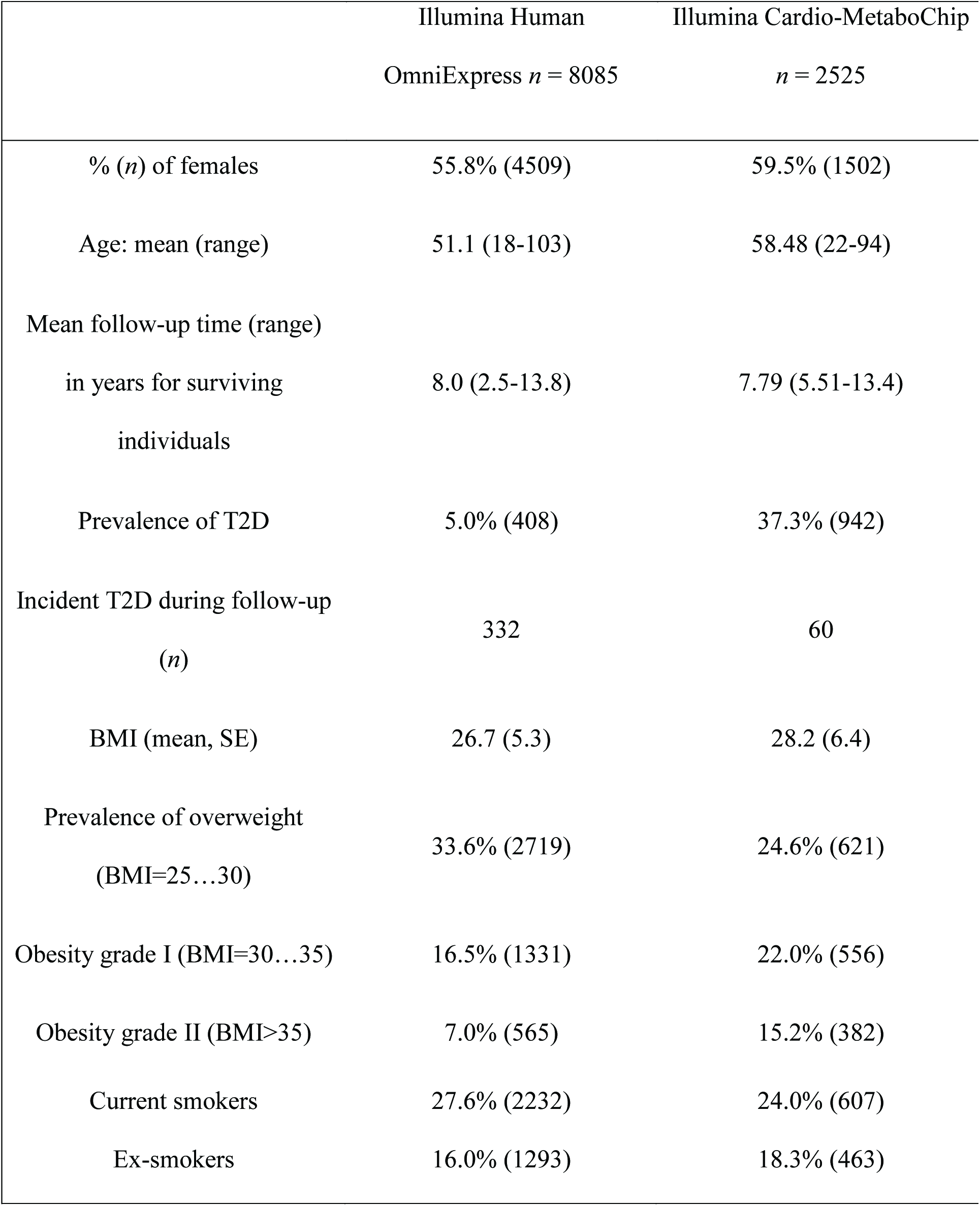
Baseline characteristics of the two genotyped subsets of the Estonian Biobank cohort. Descriptive statistics is provided separately for both subsets of Estonian Biobank cohort depending on the genotyping platform.

Genetic data used in this study have been selected as follows. First, the Estonian Cardio-Metabochip sample was used in the large-scale GWAS meta-analysis for T2D susceptibility from the DIAGRAM Consortium^11^. We therefore re-ran the meta-analysis to remove the effects of Estonian sample, as we intended to use Cardio-Metabochip for developing the optimal GRS score. Secondly, only SNPs with p-value for association with T2D less than 0.5 were taken from the meta-analysis results for further analysis. A set of independent SNPs *r^2^ ≤* 0.05 was then obtained by LD-based clumping procedure from PLINK^12^. Finally, clumped SNPs were retrieved from the Estonian Biobank database and filtered for genotyping and imputation quality and minor allele frequency, resulting in a set of 7502 SNPs for further analysis and risk score construction (Table S1). Full details about SNPs selection and weights are given in the Online Supplement.

## Statistical Analysis

Statistical analysis was performed using R version 3.1.0.^13^

### The Doubly-Weighted GRS

Let GRS_k_ denote the conventionally used GRS (also referred to as the single-weighted GRS), defined as a weighted sum of allele dosages of *k* independent SNPs, chosen on a basis of a p-value threshold from the GWAS meta-analysis (typically all independent SNPs with p-values for association less than 5*10^−8^ or 5*10^−6^). The GRS_k_ suffers from phenomena called “winners curse” – by selecting only SNPs with estimated p-values below a certain threshold, one systematically selects SNPs with effect overestimated by chance. We propose a *doubly-weighted* GRS, denoted by dGRS_k_, defined as weighted sum of all available independent SNPs, with weight for each SNP defined as a product of the GWAS parameter estimate and an estimated probability of belonging to the set of top *k* SNPs with strongest effect on the phenotype. We estimate such probability by simulating new values of potential parameter estimates based on the observed estimates and their standard errors (more details in the Supplement). We have conducted simulation studies (not presented in this paper) that demonstrated that dGRS_k_ indeed decreases the bias caused by “winners curse” in the single-weighted GRS_k_. Although in practice the algorithm requires a large number of SNPs, choosing a small value of *k* will result in near-zero weights for most of the SNPs used.

### Comparison of Different Versions of the GRS in the Association with Prevalent T2D Status

Using the data of the genotyped subsets of the Estonian Biobank cohort, we calculate both GRS_k_ and dGRS_k_ by varying *k* from 1 to all 7502 of the initially selected SNPs. The effect of each GRS is assessed using age-, sex-and genotype platform-adjusted logistic regression models for prevalent T2D status. Both BMI-adjusted and unadjusted models are fitted. The fit of (non-nested) models using a different version of the GRS as a covariate is compared using the Cox Likelihood Ratio test^14^. The GRS producing the highest log-likelihood for both BMI-adjusted and unadjusted models is selected for further validation.

The estimated T2D prevalence in individuals aged 40-79 years is visually compared across quintiles of the GRS and BMI category (< 25, 25…30, 30…35, > 35) using bar charts, while scaling the estimates to match the BMI-category-specific prevalence in the entire Estonian Biobank cohort (*n* = 28032, 2010 T2D cases in the age group 40-79). In addition, bar charts are produced to study the distribution of individuals across GRS quintiles within the subset with prevalent T2D and in the subset of obese (BMI > 35) T2D-free individuals aged 60 and older.

### Validation of the GRS in the Analysis of Incident Conditions

The GRS is further assessed for its effect on T2D incidence in individuals without prevalent T2D at baseline, all-cause and cardiovascular mortality (in all individuals), using Cox proportional hazards modelling with age as time scale. All models are adjusting for sex, smoking category (former, current) and BMI at recruitment.

The analysis was restricted to the subset of 6280 individuals aged 35-79 at recruitment (302 incident T2D cases), whereas censoring all T2D diagnoses beyond age 80, as the diagnoses in the elderly are often related to significant risk-altering co-morbidities (cancer or cardiovascular diseases). The Kaplan-Meier graph of cumulative incidence of T2D is obtained for the subset with BMI>23.

### Association of the GRS with Other Known T2D Risk Factors

The effect of GRS on BMI, Waist-Hip Ratio (WHR), plasma Glucose, Total Cholesterol, HDL-Cholesterol, LDL-Cholesterol and Triglycerides levels was estimated using age-and sex-adjusted linear regression analysis, separately in individuals with and without prevalent T2D diagnosis.

Cox Proportional Hazards model was fitted to estimate the effect of GRS on incident T2D in the subset with available glucose and lipid measurements (5373 individuals, 191 incident T2D cases), adjusting for glucose, triglyceride and HDL-cholesterol levels, as well as BMI, smoking level, sex and age at recruitment.

### Analysis of Incremental Value of GRS

For prevalent T2D, the area under the receiver operating characteristic (ROC) curve (AUC) was obtained from logistic regression fitted for individuals in age group 40-79 who were genotyped on the OmniExpress platform. For incident T2D, Harrell´s c-statistic (concordance index) from the Cox proportional hazards models for individuals aged 35-79 with no prevalent T2D diagnosis was obtained.

To study reclassification, 5-year T2D risk predictions, Cox proportional hazards models with and without accounting for GRS were fitted. Improvement in the predictions was assessed using continuous net reclassification improvement (NRI) and integrated discrimination improvement (IDI)^15^. Confidence intervals for reclassification indices and c-statistics were estimated with bootstrapping.

### Simulation Study to Investigate Heritability of the GRS

As detailed information on T2D family history is not sufficiently documented for the Estonian Biobank cohort, a simulation study is conducted to investigate possible heritability of the GRS. For random pairs of individuals (“parents”) from the genotyped Estonian Biobank subset, potential “child” genotypes are generated by combining randomly one allele from each “parental” genotype. For each simulated “child” we computed the GRS and compared that to the average GRS and to the largest GRS of the two “parents”, using Pearson coefficients of correlation and scatter plots.

## RESULTS

### Comparison of Different Versions of the GRS

Results of model fit for prevalent T2D status with GRS_k_ and dGRS_k_ for selected values of *k* are shown in Table 2. (More detailed results for values of *k* varied between 1 and 7502 are found in Table S2 and a corresponding plot of Likelihood Ratio Statistics in Figure S1). While compared to the GRS_65_ (similar to ^16^), the fit is considerably improved by using a GRS_k_ with larger number of markers – the highest log-likelihood is achieved with GRS_2100_ (BMI-unadjusted models) and GRS_600_ (BMI-adjusted models). However, when dGRS_k_ is used instead, with *k* = 300 or larger (up to 3500), the fit gets significantly better than that with any GRS_k_. The highest log-likelihood is achieved with dGRS_1400_ (BMI-unadjusted models) and dGRS_800_ (BMI-adjusted models), whereas regardless of whether the analysis is adjusted for BMI or not, dGRS_1000_ provides a fit that is not significantly different from best-fitting GRS (Cox test p-value > 0.05). **Therefore we are using dGRS_1000_ in all subsequent analyses** (weights shown in Figure S2).

**Table 2.** Summary statistics for analyses performed with different GRS versions. Estimated logistic regression parameters (Beta and SE corresponding to the effect one Standard Deviation of the GRS), Likelihood Ratio Test (LRT) statistic and the corresponding p-value for the association of different versions of the GRS with prevalent Type 2 Diabetes in the genotyped subset of the Estonian Biobank cohort. Results are reported for BMI-unadjusted and BMI-adjusted analysis. All models are adjusted for genotyping platform, age and sex.

| No BMI-adjustment |  |  |  | BMI-adjusted analysis |  |  |
| --- | --- | --- | --- | --- | --- | --- |
| GRS | Beta (SE) | LRT | p-value (LRT) | Beta (SE) | LRT | p-value (LRT) |
| GRS<sub>65</sub> | 0.34 (0.037) | 90.8 | $5.3 \times 10^{-21}$ | 0.40 (0.041) | 98.0 | $4.1 \times 10^{-23}$ |
| GRS<sub>600</sub> | 0.39 (0.037) | 113.0 | $2.1 \times 10^{-26}$ | 0.41 (0.041) | 101.4 | $7.4 \times 10^{-24}$ |
| GRS<sub>2100</sub> | 0.40 (0.037) | 121.3 | $3.2 \times 10^{-28}$ | 0.39 (0.041) | 91.9 | $9.3 \times 10^{-22}$ |
| dGRS<sub>800</sub> | 0.44 (0.037) | 151.4 | $2.3 \times 10^{-34}$ | 0.46 (0.041) | 133.0 | $9.3 \times 10^{-31}$ |
| **dGRS<sub>1000</sub>** | **0.44 (0.037)** | **153.8** | **$4.2 \times 10^{-35}$** | **0.46 (0.041)** | **132.6** | **$1.0 \times 10^{-30}$** |
| dGRS<sub>1400</sub> | 0.44 (0.037) | 154.3 | $1.8 \times 10^{-35}$ | 0.46 (0.041) | 128.5 | $8.8 \times 10^{-30}$ |

### Association of dGRS_1000_ with Prevalent T2D

The estimated Odds Ratio (OR) corresponding to one standard deviation (SD) difference in dGRS_1000_ is 1.56 (95% CI 1.45, 1.68) in the BMI-unadjusted model and 1.59 (95% CI 1.46, 1.72) in the BMI-adjusted model. The prevalence of T2D by BMI category and quintiles of dGRS_1000_ in the subset of the cohort with the age range 45-79 is shown on Figure 1 A) (see Figure S3 for a more detailed plot). Although there is no significant interaction between BMI and dGRS_1000_, the association is strongest in overweight or moderately obese individuals (25 < BMI < 35), where the number of T2D cases in the highest GRS quintile is roughly comparable to the total number of cases in the three lowest quintiles.

**Figure 1.**
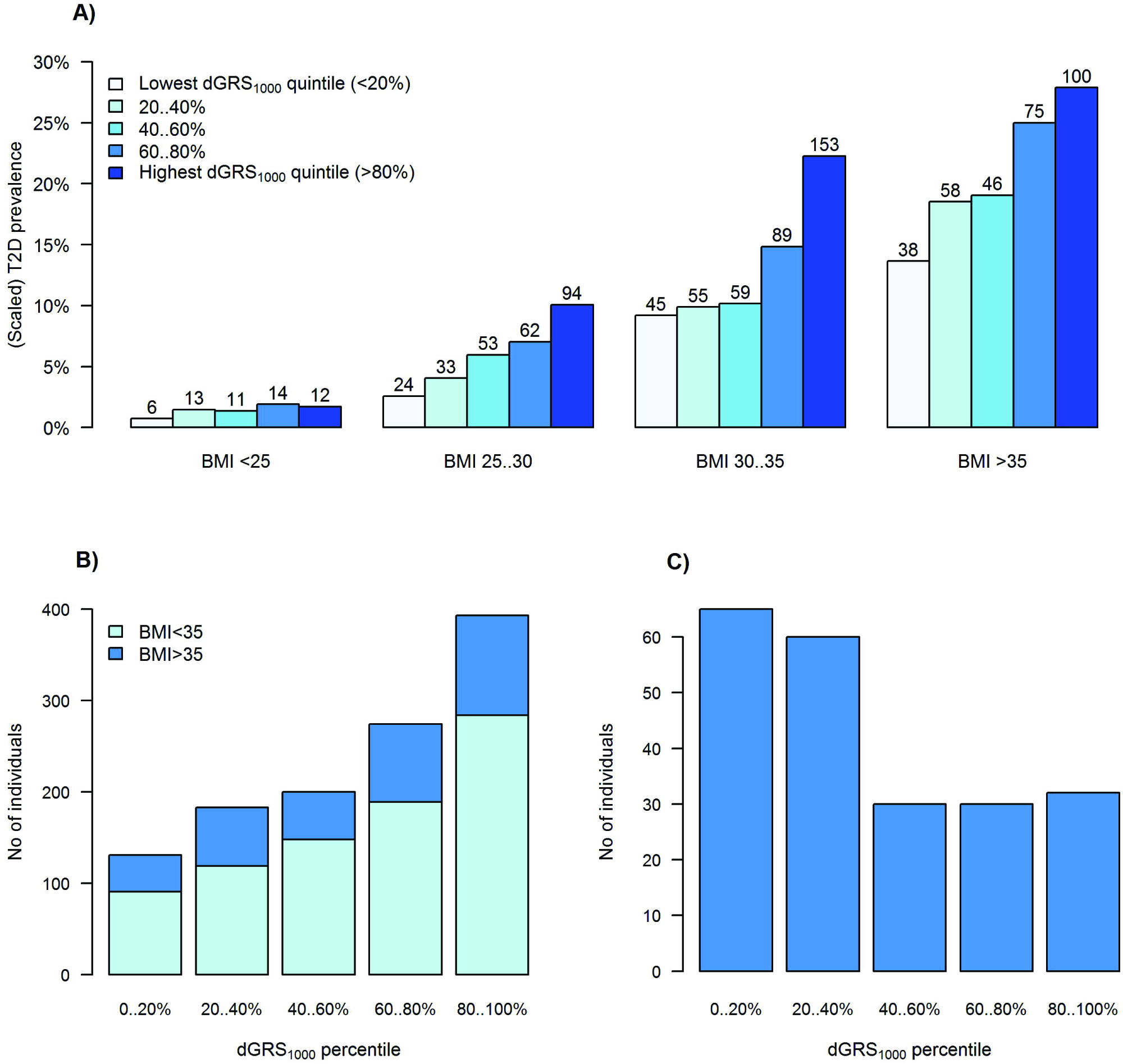
A) T2D prevalence in genotyped individuals aged 45-79 of the Estonian Biobank cohort by dGRS_1000_ quintile and BMI category. The y-axis is scaled to match the average T2D prevalence by BMI category in the entire Estonian Biobank cohort (23538 individuals aged 45-79, including 1936 cases of prevalent T2D). The total number of T2D cases in each BMI-dGRS_1000_ category is shown on the top of each bar. **B) Distribution of dGRS_1000_ in all 1181 genotyped individuals with prevalent T2D**. Distribution of dGRS_1000_ is shown among all prevalent T2D cases, with dark blue color indicating individuals with BMI > 35. **C) Distribution of dGRS_1000_ in severely obese T2D-free individuals of age 60 and over**. Distribution of dGRS_1000_ is shown among T2D-free individuals, who have BMI > 35 and are older than 60.

Figure 1 B) indicates that about one third of all prevalent T2D cases correspond to individuals in the highest GRS quintile, whereas the trend in the proportion of diseased individuals by GRS quintile is more obvious in those with BMI less than 35. On the other hand, as indicated by Figure 1 C), the majority of severely obese, but T2D-free individuals of age 60 and older, belong to the two lowest GRS quintiles.

### Validation of the GRS in the Analysis of Incident Conditions

As seen from Table 3, dGRS_1000_ has a strong effect on the hazard of developing T2D during follow-up, while accounting for age, BMI and smoking category, with more than three-fold difference in hazards between lowest and highest GRS quintile. It is also important to note that the hazard in the highest dGRS_1000_ quintile is almost two times higher than in the rest of the sample (HR = 1.90, 95%CI 1.48-2.44), indicating that this subset could be targeted for risk-reducing interventions. This is additionally supported by the fact that the highest dGRS_1000_ quintile is associated with 14% higher hazard for all-cause mortality (p = 0.019) and 27% higher hazard for cardiovascular mortality (p = 0.001).

**Table 3.** Analysis of the effect of dGRS_1000_ on incident T2D, on all-cause and cardiovascular mortality. Hazard ratios (HR) with 95% Confidence Intervals (95%CI) from Cox proportional hazards’ modelling of effect of dGRS_1000_ on incident T2D (in individuals aged 35-79 and free of T2D at recruitment) and on all-cause and cardiovascular mortality. All models are adjusted for smoking status (current or former), BMI and sex, whereas age is used as time scale.

| **Incident condition (cases/total)** |  |  |
| --- | --- | --- |
| GRS covariate | HR (95%CI) | p-value |
| **Incident T2D (302/6280), age 35-79** |  |  |
| dGRS<sub>1000</sub> (effect per 1SD) | 1.48 (1.32, 1.65) | 1.6*10<sup>-11</sup> |
| dGRS<sub>1000</sub> quintiles: 1 (< 20%) | 1 (ref) |  |
| 2 (20...40%) | 1.62 (1.04, 2.51) | 0.034 |
| 3 (40...60%) | 2.30 (1.52, 3.50) | 9.0*10<sup>-5</sup> |
| 4 (60...80%) | 2.55 (1.68, 3.88) | 1.1*10<sup>-5</sup> |
| 5 (≥ 80%) | 3.46 (2.31, 5.17) | 1.5*10<sup>-9</sup> |
| dGRS<sub>1000</sub> quintile 5 vs quintiles 1..4 | 1.90 (1.48, 2.44) | 5.3*10<sup>-7</sup> |
| **All-cause mortality (1994/10273)** |  |  |
| dGRS<sub>1000</sub> (effect per 1SD) | 1.03 (0.99, 1.08) | 0.14 |
| dGRS<sub>1000</sub> quintile 5 vs quintiles 1..4 | 1.14 (1.02, 1.27) | 0.019 |
| **Cardiovascular mortality (1069/10273)** |  |  |
| dGRS<sub>1000</sub> (effect per 1SD) | 1.06 (1.00, 1.13) | 0.046 |
| dGRS<sub>1000</sub> quintile 5 vs quintiles 1..4 | 1.27 (1.10, 1.46) | 0.0013 |

The differences in cumulative T2D incidence across dGRS_1000_ quintiles are also shown on Figure 2.

**Figure 2.**
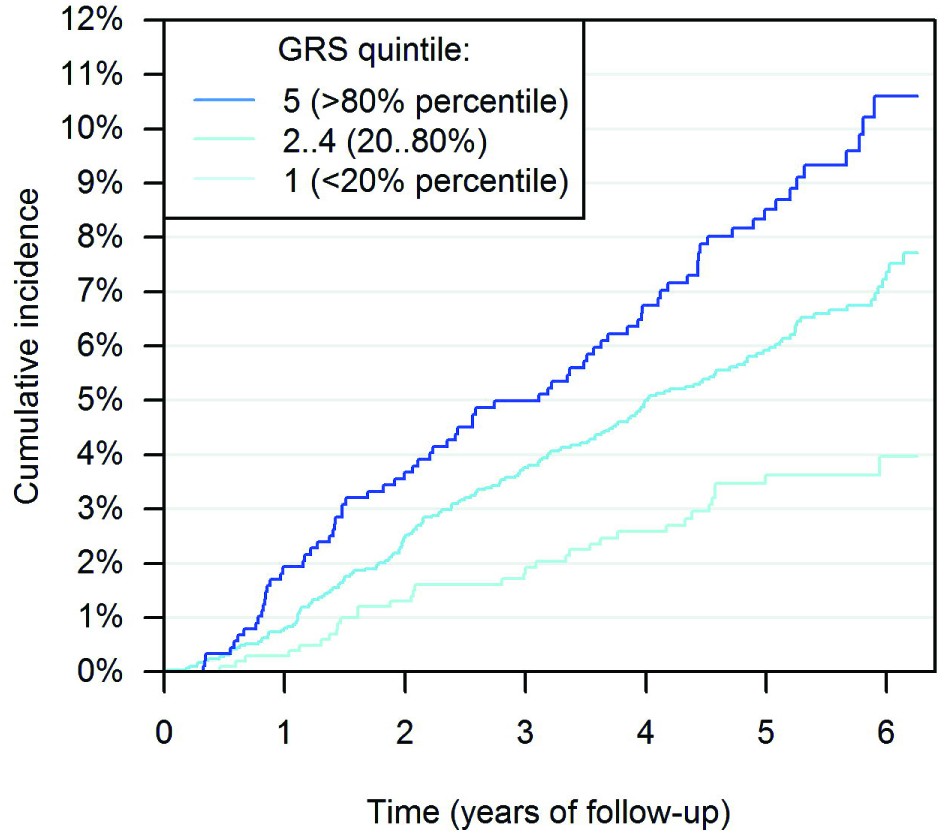
Cumulative incidence of Type 2 Diabetes in 4881 genotyped individuals free of T2D, aged 35-79 and with BMI > 23 at baseline. In the figure, 6.25-year follow-up is presented, as only 25% of individuals were followed up for more than 6.25 years. Cumulative incidence in presented separately in three dGRS_1000_ categories.

### Association of dGRS_1000_ with Known T2D Risk Factors

The estimated regression coefficients (Beta, SE, p-value) from the analysis of the effect of dGRS_1000_ on BMI, WHR, plasma glucose, total cholesterol, triglycerides, HDL-and LDL-cholesterol levels, are presented in Table S3.

In individuals without prevalent T2D, a significant positive association of dGRS_1000_ with plasma glucose *(p* = 4.7*10^−8^) and triglyceride levels *(p* = 8.8*10^−4^) and negative association with HDL-cholesterol level *(p* = 8.1*10^−5^). There is no significant association of dGRS1_00_0 with BMI found in T2D-free individuals *(p =* 0.42), but there is a weak positive association with WHR *(p =* 0.012). In individuals with prevalent T2D, a significant positive association of dGRS_1000_ with plasma glucose level *(p* = 0.0032), but no significant association with lipid profiles *(p >* 0.5) is found. The association between BMI and dGRS_1000_ in individuals with existing T2D diagnosis is found to be negative *(p* = 0.0039), indicating that individuals at high genetic risk are more likely to get T2D at a lower level of BMI than those with low genetic risk.

In the analysis of the effect of dGRS_1000_ on incident T2D, while adjusting for glucose, triglyceride and HDL-cholesterol levels, as well as BMI, smoking level, sex and age at recruitment, the HR corresponding to one SD difference in of dGRS_1000_ was estimated as 1.37 (95% CI 1.18, 1.59; *p* = 3.0*10^−5^), indicating that the effect of dGRS_1000_ has a significant effect on T2D risk that is independent of obesity and other well-known risk factors.

### Analysis of Incremental Predictive Ability of the dGRS1000

Incremental predictive ability of dGRS1_000_ was studied for both prevalent and incident T2D models. Including dGRS_1000_ in the logistic regression model for prevalent T2D improves AUC, irrespective of BMI adjustment (Figure S4). Harrell´s c-statistic increased by 0.015 (95% CI 0.006, 0.026) after adding dGRS10_00_ to the model for incident T2D. More detailed results for *c*-statistics and likelihood ratio tests are shown in the Table S4. Comparing 5-year predictions from models with and without dGRS_1000_ (Figure S5) resulted in continuous NRI of 0.264 (95% CI 0.153, 0.378). Further investigation of components of continuous NRI was undertaken as suggested^15^ - continuous NRI for events was 0.078 (95% CI −0.035, 0.187) and for non-events was 0.186 (95% CI 0.163, 0.212). IDI was 0.011 (95% CI 0.006, 0.016).

### A Simulated Example to Illustrate the Heritability of the dGRS_1000_

The Pearson correlation coefficient between offspring dGRS_1000_ and the average parental dGRS_1000_ was 0.72, whereas the correlation with the maximum of parental dGRS_1000_-s was 0.62. High average parental dGRS_1000_ does not necessarily lead to high offspring dGRS_1000_ – 33% of the children with average parental dGRS_1000_ exceeding the 80% quantile have their own dGRS_1000_ smaller than the 80% quantile (Figure S6). Also, even when one of the parents has an extremely high dGRS_1000_, it is possible that the dGRS_1000_ of the offspring is close to the median. At the same time, 15% of the children with average parental dGRS_1000_ between the 20% and 80% quantile would have their dGRS_1000_ in the highest quintile.

## DISCUSSION

We have shown a strong effect of a polygenic genetic risk score involving 7502 SNPs on the risk of T2D. The large number of SNPs included in the score is one of the main differences between our proposed GRS and earlier publications^16^, in addition to the novel weighting scheme that reduces the bias due to “winners curse”.

A large part of our work concentrates on the analysis of prevalent T2D, assessing how well the GRS discriminates between cases and non-cases. Despite the retrospective nature of this analysis - we argue that the results can be interpreted in terms of predictive power of the GRS, as SNPs cannot be affected by unmeasured confounders. The number of available prevalent cases (1181) is sufficient for acceptable power and precision in the effect estimates, as well as for efficient comparison of different versions of the GRS. However, as the main lifestyle-related predictors (BMI, physical activity and smoking level) can be affected by prevalent disease status, the analysis that is adjusted for age and sex only may be most appropriate. Our comparison of T2D prevalence across GRS quintiles and BMI categories can therefore be viewed merely as an illustration on the possible magnitude of the effect size for individuals at different body weight categories.

We propose a novel approach for the selection and weighting of SNPs in the GRS, provided a set of independent SNPs is identified. Instead of making a yes/no decision for each SNP on whether or not to include it in the score, we include a large number of SNPs, weighted by the estimated probabilities of belonging to the set of top *k* SNPs (in addition to weighting by the estimated GWAS allelic effect size). While being aware that the “winners curse” bias is not entirely removed by our procedure, we have clearly demonstrated the superiority of the proposed doubly-weighted GRS over a large variety of possible single-weighted scores.

Based on the Estonian Biobank sample, we conclude that the optimal “top” set of SNPs for T2D includes *k* = 1000 SNPs. Either decreasing *k* or increasing it (beyond 2300) would produce a GRS with weaker association with disease status. This is an indicator that it is likely that the number of independent T2D-associated genomic loci is considerably larger than currently established by most recent GWAS studies^11,17^.

One obvious issue to address is the question of independence of the GRS from other well-known risk factors for T2D, such as obesity. We have demonstrated that the effect of GRS does not depend on BMI – a similar risk level is observed for individuals at low GRS and high BMI as well as for those at relatively low BMI and high GRS level (see Figure 1 A)). Also the observed negative BMI-GRS association in diseased individuals suggests that individuals with high GRS develop the disease at lower average BMI level than those at lower genetic risk, suggesting an additive effect of the two risk factors. Despite of the fact that the GRS is associated with blood glucose, triglyceride and HDL-cholesterol levels in T2D-free individuals, the effect of dGRS_1000_ on disease incidence remains practically unchanged after adjustment for these factors.

It is well known that using c-statistics for assessing the incremental value of genetic information when strong classical predictors - such as BMI and age for T2D - are already in the model results in a very small improvement ^15^. The results of the ROC-analysis in the present study suggest that the changes in c-statistics are concordant with previously reported results^7^. However, the clinically relevant quantity is not the relative contribution of GRS in comparison with age or BMI, for instance, but its ability to distinguish between different individual risk levels in subjects of the same age and BMI. Indeed – although the overall improvement in c-statistic for the incident T2D in individuals aged 35-74 is 1.5%, this increases to 2.5% in subset with BMI in 25…35.

It has been debated whether genetic risk estimates based on DNA markers would add any meaningful information in cases where the family history of T2D is known, and it has been shown that complete family history provides better prediction than that achieved using 21 SNPs^18^. However, the highly polygenic nature of T2D suggests that parent and offspring genetic risk may actually differ considerably, as on average, only half of the risk-affecting alleles are transmitted from each parent to the child. Therefore a GRS that includes a large number of SNPs has a potential of capturing the genetic risk more accurately than family history data. Our simulation study indicates that the correlation between average parental GRS (dGRS_1000_) and child GRS is only 0.72, allowing for notable differences between the level of polygenic risk between parents and offspring. In addition, the level of environmental/lifestyle component of the risk can also differ between different family members, and therefore a high (or low) genetic risk level does not necessarily result in occurrence (or non-occurrence) of the disease. Moreover, in current family structures, it is increasingly difficult to get detailed information, if any, from both parents. Therefore, our study suggests that as the number of SNPs included in the GRS increases, the accuracy of GRS-based risk estimate improves in comparison to that based on family history.

Trans-ethnic analysis of T2D have suggested that the effect sized for common SNPs are relatively homogenous across ethnicities ^17^. However, as the effect sizes in the GWAS meta-analysis used in our study are calculated based on cohorts of predominantly European descent, one should be still cautious about extrapolating our results to other populations without further validation. We recommend using population-based biobank data, where available, to validate the GRS in each target population before implementing it in the actual risk prediction.

Our study also indicates that for diseases with polygenic nature, a very large discovery sample is needed for the GWAS (meta-analysis) to provide effect estimates as a basis of a GRS. Despite the fact that the estimates used in our study are based on a very large sample size (34840 T2D cases and 114981 controls), using an even larger sample could further improve the predictive accuracy of a GRS.

In addition, we also see room for further methodological improvements, by further reducing the “winners curse” bias and/or allowing for possible inclusion of correlated SNPs.

In conclusion, the doubly-weighted GRS computed on the basis of a large number of SNPs leads to improvement in predictive ability compared to versions of the GRS that include SNPs with established genome-wide significance only and/or use a different weighting scheme. As individuals in the highest quintile of the dGRS_1000_ were observed to have approximately 4 times higher hazard of developing T2D before the age of 70, the effect size is clinically meaningful. The proposed GRS works both for long-term predictions (from birth on), but also in the short term, when other risk factors are already accounted for. Our results indicate that a GRS with high accuracy, such as dGRS_1000_, would significantly improve the best existing risk assessment algorithms for T2D, encouraging its implementation in the practice of personalized medicine.

## Acknowledgments

### Contributions

KF, KL, RM, AMo and AMe were involved in planning the study design. KL, AMo and RM contributed to data management. KF and KL analysed and interpreted the data. KF and KL wrote the first draft of the manuscript. All co-authors - KF, KL, RM, AMo, and AMe - read the manuscript and contributed to the final version.

### Disclosure

The authors declare no conflict of interest.

### Funding

EGCUT received financing from Estonian Research Council Grant GP1GV9353 and IUT20-60, Center of Excellence in Genomics (EXCEGEN), University of Tartu (SP1GVARENG) and EU structural Fund through Archimedes Foundation, grant no. 3.2.1001.11-0033.

